# Therapeutically targetable Th17-derived miR-721 drives autoimmune myocarditis through PPARγ repression

**DOI:** 10.64898/2026.03.24.713340

**Authors:** Ignacio Ruiz-Fernández, Raquel Sánchez-Díaz, Rafael Blanco-Domínguez, Enrique Ortega-Sollero, Ruth Ortego-Moltó, Daniela Quiroga-Ortiz, Hortensia de la Fuente, José Martínez-González, Luis Jesús Jiménez-Borreguero, Beatriz López-Melgar, Fernando Rivero, Fernando Alfonso, Francisco Sánchez-Madrid, Mercedes Ricote, Pilar Martín

**Affiliations:** Centro Nacional de Investigaciones Cardiovasculares (CNIC), Madrid, Spain; Centro de Investigación Biomédica en Red de Enfermedades Cardiovasculares (CIBER CV), Madrid, Spain; Escuela de Doctorado. Universidad Autónoma de Madrid, Spain; Gulbenkian Institute for Molecular Medicine, Lisbon, Portugal; Department of Immunology, Instituto de Investigación Sanitaria Hospital Universitario de La Princesa (IIS-Princesa), Universidad Autónoma de Madrid (UAM), 28006 Madrid, Spain; Instituto de Investigación Biomédica Sant Pau (IIB-Sant Pau), Barcelona, Spain; Instituto de Investigaciones Biomédicas de Barcelona-Consejo Superior de Investigaciones Científicas (IIBB-CSIC), Barcelona, Spain; Department of Cardiology, Instituto de Investigación Sanitaria Hospital Universitario de La Princesa (IIS-Princesa), 28006 Madrid, Spain; Centro Nacional de Biotecnología (CNB), Consejo Superior de Investigaciones Científicas (CSIC), Madrid, Spain

**Keywords:** Autoimmune Myocarditis, Th17 cells, miR-721, PPARγ

## Abstract

**BACKGROUND:** Myocarditis is an inflammatory cardiac disease in which Th17-driven immune responses contribute to progression toward dilated cardiomyopathy and heart failure. Current therapies mainly rely on corticosteroids but lack specificity, while the role of miR-721, synthesized by Th17 cells, remains largely unexplored in disease pathogenesis.

**METHODS:** We characterized the presence of mmu-miR-721 and its human homolog hsa-RNA-Chr8:96 in extracellular vesicles (EVs) secreted by Th17 cells from IL-17eGFP mice with experimental autoimmune myocarditis (EAM) and myocarditis patients. MxCre-Pparg^fl/fl^ mice and luciferase reporter assays were used to validate the target genes of miR-721 and hsa-RNA-Chr8:96, respectively. The functional role of miR-721 in EAM was investigated by lentiviral vectors overexpression and inhibition using miRNA sponge molecules. Th17 responses and heart inflammation were assessed and echocardiography was performed after in vivo blockade of mmu-miR-721 in EAM mice.

**RESULTS:** Both mmu-miR-721 and hsa-RNA-Chr8:96 were encapsulated in EVs and secreted by Th17 cells of mice and patients with myocarditis. Overexpression of mmu-miR-721 in draining-lymph node cells from EAM mice inhibited *Pparg* transcription, leading to increased RORγt and IL-17 expression and promoting Th17 differentiation. In contrast, in the absence of *Pparg*, a target of miR-721, no differences in RORγt expression were observed, indicating that miR-721 promotes Th17 responses through repression of *Pparg*. Human *PPARG* was validated as a target gene of hsa-RNA-Chr8:96 and its overexpression in peripheral blood leukocytes downregulated *PPARG* mRNA levels, suggesting similar pathways involved in human pathology. In vivo blockade of mmu-miR-721 increased *Pparg* expression, reducing RORγt and IL-17 activation in T cells and leading to decreased leukocyte infiltration in the heart and improved cardiac function.

**CONCLUSIONS:** miR-721 is released by Th17 cells in EVs and promotes Th17 responses during myocarditis through repression of PPARγ, identifying this miRNA as both a mechanistic driver of disease and a potential therapeutic target.

**Graphical abstract:** 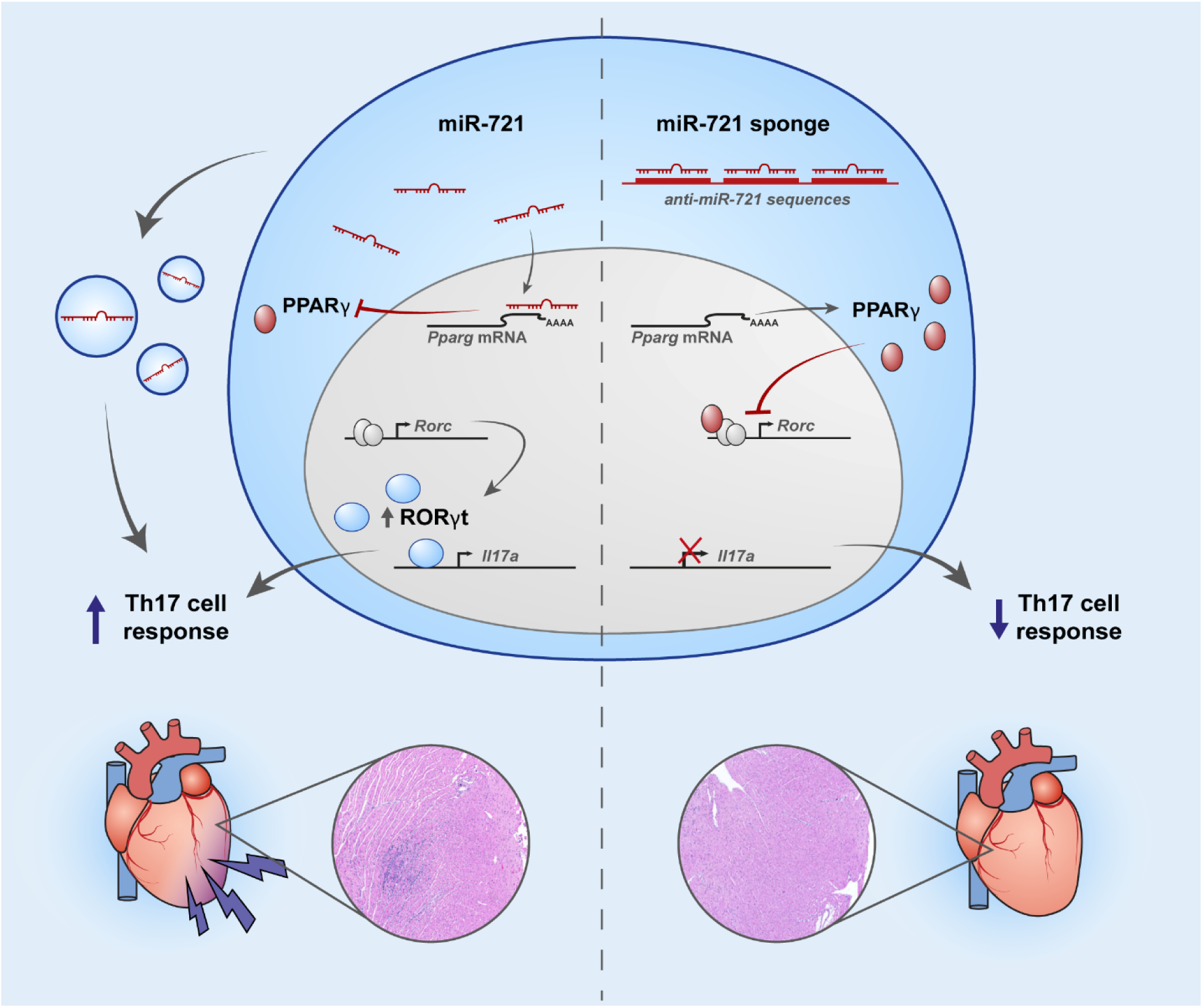

**Novelty and significance:** *What is known?:*

- miR-721 and its human homolog are upregulated in the plasma of mice and humans with myocarditis
- Th17 cells synthesize miR-721
- Mmu-miR-721 targets *Pparg* mRNA

*What new information does this article contribute?:*

- miR-721 is sorted into extracellular vesicles in the context of acute myocarditis
- miR-721 enhances Th17 differentiation via the *Pparg*/*Rorc* double inhibitory axis.
- Hsa-RNA-Chr8:96 targets human *PPARG* mRNA for degradation, inhibiting its expression
- Blockade of miR-721 dampens acute myocarditis development *in vivo* This study reveals a novel miRNA-based therapeutic strategy to inhibit Th17 responses and treat myocarditis. Using the experimental autoimmune myocarditis model, the authors unravel the mechanisms by which mmu-miR-721 can enhance Th17 responses and show how targeting this regulatory molecule could ameliorate the progression of the disease. Remarkably, this regulatory axis is suggested to be present in humans as well, since *PPARG* gene is validated as a target gene for hsa-RNA-Chr8:96. These findings highlights the potential of miR-721 not only as a diagnostic tool but also as a cell-specific therapeutic target to control Th17 responses in the clinical setting.

## Introduction

Myocarditis is a clinically challenging entity characterized by a generalized inflammation of the cardiac muscle originated by infectious or sterile stimuli^1^. Sustained inflammation may progress towards irreversible dilated cardiomyopathy and heart failure. Nowadays, management of myocarditis patients typically consists of unspecific treatments to decrease inflammatory damage and heart dysfunction^2^. During myocarditis, the activation of autoreactive T helper 17 (Th17) cells, which secrete interleukin-17 (IL-17), orchestrates the immune response and is considered as a key event of the progression towards chronic scenarios regardless myocarditis etiology^3^. Therefore, specific therapeutic strategies targeting this cell type would be of great help to improve the outcome of myocarditis patients.

Recently, we identified that mmu-miR-721 and a human equivalent, hsa-RNA-Chr8:96, are synthesized by Th17 cells in the context of murine or human myocarditis, respectively, acting as efficient circulating biomarkers of this pathology^4^. However, it remains unknown whether the miRNA produced by Th17 cells is released as a free molecule or packaged within extracellular vesicles (EVs). Some miRNAs can be sorted into extracellular vesicles (EVs) and shuttled in a more protected manner to their target cells. EV-derived miRNAs offer higher detection sensitivity and specificity compared to traditional serum biomarkers for liquid biopsy^5^. Consequently, knowing whether miR-721 is sorted and secreted in EVs is important for better understanding its potential as a biomarker and crucial for elucidating the role and targets of this miRNA in myocarditis pathology.

In the previous study, miR-721 biological pathways and putative target genes were analyzed, revealing its association with T cell homeostasis and differentiation^4^. Specifically, targets related to Th17 cell differentiation were identified. Among these target genes, *Pparg* was a notable validated target of miR-721^6^ with regulatory roles in cardiovascular pathophysiology^7^. In this regard, PPARγ has been identified as a negative regulator of Th17 differentiation in both humans and mice^8^. Th17 differentiation requires RORγt expression and is induced by TGF-β/IL-6 or IL-21. Activation of PPARγ in CD4^+^ T cells selectively suppressed Th17 differentiation by inhibiting RORγt expression. Pharmacologic activation of PPARγ prevented *Rorc* transcription by maintaining corepressor binding to its promoter. PPARγ’s role was confirmed in T cell-specific knockout models and in human CD4^+^ T cells from healthy controls and MS patients. Thus, PPARγ is a promising target for treating Th17-mediated autoimmune diseases^8^. Moreover, PPARγ ligands ameliorated Experimental Autoimmune Myocarditis (EAM) by inhibiting the activation of self-sensitive T cells^9^. Together, these experimental evidence suggests that targeting PPARγ may represent a strategy to reduce Th17 responses in myocarditis.

In the present work, we describe the presence of mmu-miR-721 and hsa-RNA-Chr8:96 in plasma EVs from EAM and myocarditis patients, respectively. Mmu-miR-721 overexpression in draining lymph nodes (dLN)-derived cells enhances Th17 cell responses in EAM. Mechanistically, mmu-miR-721 directly targets and downregulates *Pparg* mRNA, which results in de-repression of RORγt expression and Th17 cell expansion. *PPARG* was validated as a target gene for hsa-RNA-Chr8:96, suggesting that similar pathways are controlled in the human pathology. Conversely, *in vivo* blocking of mmu-miR-721 with anti-miR sponge molecules^10^ during EAM results in diminished numbers of Th17 cells in the heart and, therefore, reduced cardiac inflammation and dysfunction, leading to the overall protection from myocarditis progression. In conclusion, these findings identify miR-721 inhibition as a potential strategy to specifically restrain pathogenic Th17 responses in myocarditis.

## Methods

### Mice

Experimental groups were matched for sex and age using littermate controls on either C57BL/6 or BALB/c backgrounds, depending on the experimental model and strain susceptibility. C57BL/6 Foxp3-mRFP/IL17-eGFP doble reporter mice were kindly provided by Dr. R. A. Flavell (Yale University). Mx^WT/*Cre*^ *Pparg^fl/fl^* C57BL/6 mice were used, and Cre recombinase expression was induced by intraperitoneal administration of 300 µg polyinosine-polycytosine (pIpC) on days 0, 2, and 4. Efficient *Pparg* deletion was confirmed from day 10 onwards. Animals were housed and used under specific-pathogen-free (SPF) conditions at the Centro Nacional de Investigaciones Cardiovasculares (CNIC) animal facility. All animal procedures were approved by the ethics committee of the Comunidad Autónoma de Madrid (PROEX 17.5/25; in force) and conducted in accordance with the institutional guidelines that comply with European Institutes of Health directives (European Institutes of Health. 2010).

### Mouse model of myocarditis

For EAM induction, 8-12 weeks-old BALB/c mice were immunized subcutaneously on days 0 and 7 with 100 µg/0.2 mL MyHC-α peptide (MyHCα_614–629_) Ac-RSLKLMATLFSTYASADR-OH emulsified 1:1 in complete Freund adjuvant (CFA, 1 mg/mL) and PBS. For inducing EAM in non-susceptible strains, 150 ng pertussis toxin from *Bordetella pertussis* was administered intraperitoneally at day 0 and 2 to ensure disease development. When indicated, the peptide was replaced with PBS as a control. Acute phase of disease starts from day 14 and EAM is evaluated at day 21, when tissues were harvested and plasma samples were extracted.

### Mouse models of myocardial infarction

For MI induction, a permanent ligation of the left anterior descending (LAD) coronary artery was performed. Briefly, mice were anesthetized with sevoflurane (4%) and intubated using a 24-gauge catheter with a blunt end. Mice were artificially ventilated with a mixture of O_2_ and air (1:1, vol:vol) using a rodent ventilator (*Minivent 845*, Harvard) with 160 strokes/min in a total volume of 250μl. Mice were placed on a heating pad to maintain body temperature at 37°C. A thoracotomy was performed through the fourth left intercostal space, the pericardium was opened and the heart was exposed. The proximal LAD coronary artery was localized and permanently ligated by passing a 7-0 silk suture around the artery. Sham operated mice were analyzed in parallel as controls of surgery when indicated. Peripheral blood was collected from the tail vein in EDTA-tubes.

### *In vivo* blocking of mmu-miR-721

For the blocking of mmu-miR-721 expression *in vivo* we use a Sponge single-stranded molecule consisting of 4 tandem-repeated reverse complementary (RC) mature miR-721 sequence, separated by three 4-6 bp-long spacer sequences as follows: (5′ggccgc**ttcccccttttaattgcactg**cgat**ttcccccttttaattgcactg**accggt**ttcccccttttaattgcactg**tcac**ttccccct tttaattgcactg**c 3′). A scramble sequence encoding a fragment of a recombinant GFP of similar size was used as control. mmu-miR-721 RC sequence: ttcccccttttaattgcactg. Scramble sequence: agaagaacggcatcaaggtga. In order to inhibit miR-721 *in vivo* we administered 0.2 nmol of the sponge or scramble molecule emulsified with the CFA plus MyHCα-peptide subcutaneously at the second immunization (day 7). EAM development was evaluated at day 21 and the inhibition efficiency of miR-721 by qPCR was assessed in four animals per group, which were sacrificed and dLNs were collected 2 days after administration of the sponge/scramble molecules.

### Echocardiography acquisition

Transthoracic echocardiography was performed by a blinded expert operator using a high-frequency ultrasound system (Vevo 2100, *Visualsonics Inc*., Canada) with a 30-MHz linear probe. Two-dimensional (2D) and M-mode (MM) echography were performed at a frame rate over 230 frames/sec. Mice were lightly anesthetized with 0.5-2% isoflurane in oxygen, adjusting the isoflurane delivery to maintain the heart rate in 450±50 bpm. Mice were placed in supine position and maintained normothermia using a heating platform and warmed ultrasound gel. A base apex electrocardiogram (ECG) was continuously monitored. Longitudinal and short axis views of the left ventricle were acquired at the papillary muscles level for M-mode and also medium and apical levels for 2D. From short-axis and long-axis views, end-systolic and end-diastolic LV volumes, mass and ejection fraction (LVEF) were calculated using the area-length method (Vevo 2100 Workstation software).

For the analysis of the wall motion score using echocardiography, regional left ventricular function was evaluated in the parasternal long-axis view. The left-ventricular wall was subdivided into six segments (basal, mid, and apical in the anterior and posterior walls). Each segment was scored by an independent blinded evaluator based on its motion and systolic thickening, according to the guidelines of the American Society of Echocardiography^11^: (1) normal or hyperkinetic, (2) hypokinetic (reduced thickening), (3) akinetic (absent or negligible thickening, e.g., scar), and (4) dyskinetic (systolic thinning or stretching, e.g., aneurysm). The number of dysfunctional segments was quantified, and the total wall motion score index (WMSI) representing the sum of the score of the six individual segments in each heart was calculated.

### Histology

Hearts were fixed in 10% buffered formalin and were sliced and stained with hematoxylin and eosin (HE) to observe infiltrating leukocytes and assess inflammation. The percentage of immune cell infiltration was determined as the percentage of cell infiltration compared to the overall size of the heart section using a microscope eyepiece grid and Image J Program (Rasband, W.S., ImageJ, U.S. National Institutes of Health, Bethesda, Maryland, USA, https://imagej.nih.gov/ij/, 1997-2018. Schneider, C.A., Rasband, W.S., Eliceiri, K.W).

### Collection of human blood samples

MiRNA expression was analyzed by qPCR in plasma and EV-plasma compartment from acute myocarditis patients (n=32)), ST-segment elevation myocardial infarction (STEMI) patients (n= 35), NSTEMI (non-STEMI) patients (n=24) and healthy volunteers (n= 25) (**Supplementary Table 1**).

Blood samples were collected within the first were collected within the first 24h after admission in BD Vacutainer-EDTA tubes (*BD Plymouth*, UK). Samples were processed up to 24 h from collection and during this time they were kept at 4°C. Plasma samples were obtained by centrifugation at 2000g at 4°C, aliquoted and stored at −80°C until total RNA extraction or PEG precipitation.

### Isolation of draining lymph node-derived cells

Immune cells were obtained from dLN (inguinal, axillary and brachial) from 8-12 weeks-old BALB/c mice immunized subcutaneously with 100 µg/0.2 mL MyHC-α peptide (MyHCα_614–629_) emulsified 1:1 in CFA (1mg/mL) and PBS, and 150 ng pertussis toxin from *Bordetella pertussis* was administered intraperitoneally at day 0 and 2 for C57BL/6 Foxp3-RFP/IL17A-eGFP mice. by placing the tissue into a 70-μm cell strainer (*BD Falcon*) and using the plunger end of the syringe the tissue was mashed through the cell strainer into a petri dish. The cell suspensions were filtered through the cell strainers and washed with PBS+0.05% BSA+0.01% EDTA buffer.

### Isolation of heart-infiltrating cells

Heart was perfused with 10 ml of ice-cold PBS and extracted from the chest cavity. Then, hearts were minced and digested with collagenase IV (100 U/mL; *Gibco*) for 45 minutes at 37°C in constant shaking. The resultant cell suspensions were filtered through 40-μm cell strainers (*BD Falcon*) and washed twice with PBS+0.05% BSA+0.01% EDTA buffer. Erythrocytes were removed using a hypotonic buffer. Leukocyte number was evaluated by microscopy and normalized per heart weight.

### Flow cytometry

Single cell suspensions of mouse lymph nodes or heart infiltrating leukocytes were incubated in PBS +0.05% BSA +0.01% EDTA buffer with fluorochrome-conjugated antibodies. Cells were cultured overnight for bystander activation in plates coated with 2μg/ml of purified anti-CD3 (145-2C11 clone, *BD PharMingen*) in complete RPMI medium (*Gibco*) with 1μg/mL of α-CD28 (clone 37.51, *BD*) before cell staining. For cytokine production assessment, cells were stimulated with 50 ng/ml of phorbol myristate acetate (PMA, *Sigma Aldrich*), 1 μg/ml of ionomycin (*Sigma Aldrich*) and 1 µg/ml of Brefeldin A (*Sigma-Aldrich*) in complete culture medium for 4 additional hours. Staining of membrane markers was performed by incubating single cell suspensions with fluorochrome-conjugated antibodies in PBS +0.05% BSA +0.01% EDTA for 15 minutes at 4 °C. Antibodies targeting the following markers were used: CD3 (BD, clone 145-2C11), CD4 (BD, clone RM4-5), CD11b (*BD*, clone M1/70), CD45.2 (*Biolegend*, clone 104), F4/80 (*Biolegend*, clone BM8). When required, membrane-stained cells were subsequently fixed and stained with the following intracellular or intranuclear protocols:

For RORγt evaluation, nuclear staining was performed using the Foxp3/Staining Buffer kit (*eBioscience*), following the providers’ instructions. Anti-RORγt (BD, clone Q31-378) was used.

For effector T helper cell evaluation, cells were fixed with PBS 2% paraformaldehyde for 10 minutes at room temperature and intracellularly stained with conjugated-antibodies against IL-17A (BD, clone TC11-18H10) in PBS +0.05% BSA +0.01% EDTA +0.5% saponin for 45 minutes at room temperature.

Cells were analyzed in a LSRFortessa or FACSymphony Flow Cytometer and the data were processed with FlowJo v10.0.4 (Tree Star). When indicated, Foxp3-RFP/IL17A-eGFP mice were used to sort Foxp3+ and/or IL-17+ cells in a BD FACSAria II cell sorter. For EV characterization by flow cytometry, Novocyte Opteon Spectral Flow Cytometer and NovoExpress software (Opteon) was used.

### Quantification of miRNA and mRNA expression by quantitative PCR

RNA was extracted with miRVANA miRNA Isolation kit (*Invitrogen*) or miRNeasy mini kit (*Qiagen*) as recommended by the manufacturer. For plasma samples, the former kit was used with slight modifications. Briefly, 600µl of the Lysis/Binding buffer (mirVana miRNA Isolation Kit, *Invitrogen*) were added to 200 µl of human /50-100 µl of mouse plasma.

For miRNA detection, the same volume of RNA solution of each sample was used for the RT reaction in plasma samples and RT was performed from 100 ng of total RNA for cell-derived RNA. Reverse transcription was performed using the miRCURY LNA RT kit (*Qiagen*) and microRNA expression was analyzed by real-time quantitative PCR using specific microRNA LNA PCR primer sets (*Qiagen*) for each microRNA and the miRCURY LNA SYBR Green PCR Kit (*Qiagen*). RNA Spike-in kit (*Qiagen*) was used as an exogenous control of RNA extraction according to the manufactureŕs instructions. Two synthetic RNA spike-ins in different concentrations were used to control the yield of RNA extraction (UniSp5/UniSp2). Synthetic UniSp6 was used to monitor RT-PCR yield. The use of these spike-ins as exogenous controls allowed us to exclude samples not meeting amplification standards. Normalization was performed versus the spike-ins controls using the 2^−ΔCt^ method. MiR-423-3p, miR-103a-3p and let-7a-5p were proposed by GeNorm and NormFinder as the most stable miRNAs among groups of samples and were used as endogenous normalizers.

For mRNA quantification, reverse transcription was performed with DNAse-treated RNA using the High-Capacity cDNA RT Kit (*Applied Biosystems*). Then, gene expressions were measured by real-time qPCR using SYBR Green PCR Mix (*Applied Biosystems*) and mRNA-specific primers (*Thermo Fisher Scientific*). Real-time qPCR analyses were performed with an QuantStudio™ 5 Real-Time PCR System, 384-well (*Applied Biosystems*). *Gapdh* and *Actb* genes were used as endogenous normalizers. Relative gene expression was determined using the 2 ^−ΔΔCt^ method.

### Isolation and characterization of extracellular vesicles

For dLN-EAM induction, 8-12 weeks-old Foxp3-RFP/IL17A-eGFP mice were immunized subcutaneously with 100 µg/0.2 mL MyHC-α peptide (MyHCα_614–629_) emulsified 1:1 in CFA (1mg/mL) and PBS, and 150 ng pertussis toxin from *Bordetella pertussis* was administered intraperitoneally at day 0 and 2. After 6 days, dLN-EAM cell suspensions were cultured (2×10^7^ cells/ml) during 48h in plates coated with 2μg/ml of purified anti-CD3 (145-2C11 clone, *BD PharMingen*) using exosome-free TexMACS media (*Miltenyl Biotec*) in the presence of MyHC-α peptide (10 µg/ml), 1μg/mL of α-CD28 (clone 37.51, *BD*) and IL-23 (10 ng/ml, *R&D*). For characterization of Th17^EAM^ and Treg^EAM^-derived EVs, cells were sorted using a BD FACs Aria II cell sorter or iCyt Synergy 4L cell sorter for detection of mRFP and EmGFP after staining of CD4 as mentioned above. Resulting cells were cultured in plates coated with 2μg/ml of purified anti-CD3 (145-2C11 clone, *BD PharMingen*) in TexMACS media for 48h with MyHCα peptide and either IL-2 (2 ng/ml, *R&D*) for Foxp3^mRFP+^ cells or IL-23 (10 ng/ml *R&D*) for IL-17^eGFP+^ cells. The resulting supernatants (SN) were destined for characterization of EVs or analysis of miRNA levels.

For EVs isolation, Polyethylene glycol (PEG) was employed as previously described^12^ from mouse/human plasma or supernatant from the above cultures. Briefly, cell debris were pelleted by centrifugation at 1000g for 30 min. EVs were precipitated by adding 0.4 volumes of 50% PEG 6000 (Sigma Aldrich) in 375 mM NaCl followed by 30 min of incubation at 4°C. Samples were then spin down at 1500g for 30 min at 4°C. For the characterization of EVs derived from immune cells, culture-SN EVs were analyzed by Nanosight LM10 and NTA 2.3 Software (NanoSight, Wiltshire, UK) and Zeta Sizer (ZS90 Malvern Instruments) and Novocyte Opteon Spectral Flow Cytometer after removal of cell debris by spinning down at 10000g for 30 minutes.

### Evaluation of miRNA secretion into EVs

The ratio EVs/Total-Supernatant and EVs/Total-plasma was calculated as follows. For assessement of the levels of miRNAs, the volume of liquid (200 µl SN or plasma) was split in half; after PEG-isolation, EVs from SN were solubilized in TexMACs and EVs from plasma were solubilized in saline buffer, (1:1 volume). Six volumes of Lysis/Binding buffer (mirVana miRNA Isolation Kit, *Invitrogen*) were added, plus the number of spike-ins controls recommended by the manufacturer (RNA Spike-in kit, *Exiqon*). RNA extraction was then performed as described above. Without adjusting concentration, the same volume of RNA solution of each sample was used for RT-qPCR. Normalization was performed versus the spike-ins controls using the 2^−ΔCt^ method to obtain linear values of relative quantity and avoid differences in the yield of the protocol. The ratios were then calculated using linear quantities as described^12^. The maximum value 1 indicates that the specific microRNA detected in plasma/SN samples is fully encapsulated into EVs; lower values indicate that part of this microRNA is released free in the medium and is lost after EV precipitation.

### Lentiviral vectors generation

For mmu-miR-721 overexpression using lentiviral vectors, a plasmid containing the pre-miRNA, the product of nuclear processing by the Drosha-DGCR8 complex, was generated. Mature miRNA sequence flanked by approximately 100 bp was cloned into a pHRSIN/SINBX expression plasmid (*Addgene*). For human homolog miRNA overexpression, pre-hsa-RNA-Chr8:96^4^ was cloned into a pCDH-CMV-MCS-EF1α-copGFP expression construct (*System Bioscience*). Lentiviral particles were generated by co-transfection of the plasmid of interest with or without the miRNA encoding region, pCMV-dR8.91 (*Addgene*) and pMD2G (*Addgene*) plasmids, using the calcium phosphate method in HEK-293 cells. Lentiviral vectors were titrated by transduction of Jurkat ATCC growing in exponential phase and analysis of *mmu-miR-721* or EmGFP by qPCR and FACS, respectively.

### Overexpression of mmu-miR-721 in dLN^EAM^ cells

dLN-derived cells were obtained 6 days after EAM induction of WT and Mx^WT/*Cre*^ *Pparg^fl/fl^* mice. The MOI 1, MOI 2 and MOI 5 doses were tested to infect 600,000 cells. After extensive washing, cells were cultured over 48h for bystander activation in plates coated with 2μg/ml of purified anti-CD3 (145-2C11 clone, *BD PharMingen*) in complete RPMI medium (*Gibco*) with MyHCα-p (10 µg/ml), 1μg/mL of α-CD28 (clone 37.51, *BD*) and IL-23 (10 ng/ml, *R&D*). A fraction of the cells was used to assess overexpression of miR-721 and expression of *Rorc*, *Il17a* and *Pparg* by qPCR. For cytokine production assessment by FACS, cells were stimulated with 50 ng/ml of phorbol myristate acetate (PMA, *Sigma Aldrich*), 1 μg/ml of ionomycin (*Sigma Aldrich*) and 1 µg/ml of Brefeldin A (*Sigma-Aldrich*) in complete culture medium for 4 additional hours.

### Overexpression of mmu-miR-721 in primary human peripheral leukocytes

Peripheral blood leukocytes (PBLs) were isolated from blood samples using Ficoll-Isopaque (density=1.121 g/ml) gradient centrifugation and subsequently transduced with the generated lentiviral vector and control (MOI 5). After washing, cells were cultured over 48h for bystander activation in plates coated with 2μg/ml of purified anti-CD3 (OKT3 clone, *BD PharMingen*) in complete RPMI medium (*Gibco*) with 1μg/mL of α-CD28 (clone L293, *BD Biosciences*). Expression levels were then analyzed by qPCR.

### Luminescence assay

HEK-293 cells (1 × 10^5^) were co-transfected with a psiCHECK2 construct (*Promega*) containing *PPARG* 3’UTR region cloned downstream hRluc (Renilla luciferase) and 0.5 μg of pCDH-CMV-MCS-EF1α-copGFP expression construct (*System Bioscience*) with or without hsa-RNA-Chr8:96-encoding sequence. After 24 h, cells lysates were analysed for luciferase activity using Dual-Glo Luciferase Assay System (*Promega*). The values of Renilla luciferase (RL) were normalised by firefly luciferase (RL) and by the values obtained with cells transfected with the empty construct.

### Statistical analysis

For statistical analysis, the normality of the distributions was first evaluated using Shapiro-Wilk’s test for mouse experiments and D’Agostino-Pearson’s test for human data with higher sample size. When distributions are normal, unpaired Student’s *t*-test was used to compare only two groups of samples and one-way ANOVA analysis with Holm Sidak’s *post hoc* test when more than two groups were compared. If distributions were non-normal, Mann-Whitney’s U-test was used for the analysis of two groups and Kruskal-Wallis test with Dunn’s *post hoc* test was used for multiple comparisons. Data analyses were performed with *GraphPad Prism 10.4* software.

## Results

### Mmu-miR-721 and hsa-RNA-Chr8:96 are secreted by Th17 cells into extracellular vesicles

During EAM, mmu-miR-721 is synthesized by Th17 cells^4^; however, it remains undetermined whether this miRNA is shuttled into the circulation as a free molecule or preferentially packaged within extracellular vesicles (EVs). To address this, we analyzed the presence and distribution of mmu-miR-721 and its human homolog in culture supernatants and plasma, comparing total and EV-enriched fractions after PEG-based isolation and assessing their relative abundance in each compartment. Circulating miRNA species in plasma are protected from ribonuclease degradation by encapsulation into EVs and the sequence of mature miRNAs have been postulated to predict their sorting into EVs, especially guanine-rich sequences^13^. The 3′ end positions of mmu-miR-721 and mmu-miR-483-5p sequences present the EXOmotif *GGNG,* in which the N can be equally A, C or G indistinctly^14^ (**Figure 1A**). To determine the potential of several miRNAs previously described in cardiovascular disease^4^ to be exported into EVs, we cultured dLN-EAM cells, 6 days after immunization with MyHCα-p, for 48h and isolated EVs from supernatants (SN) by precipitation with polyethylene glycol (PEG)^12^ (**Figure 1B** and described in Methods). Both mmu-miR-721 and miR-483-5p were shown preferentially secreted into EVs as their ratios of relative quantity between EVs and total SN compartment can be considered = 1, unlike other miRNAs released free into the SN whose EV/total SN ratio is < 1 (**Figure 1C and S1A-B**). To further assess the contribution of Th17 and Treg compartments, dLN-EAM Th17 and Treg cells were sorted and cultured for 48h (**Figure 1B and S1C**). The analysis of miRNAs in the EVs of sorted Th17^EAM^ and Treg^EAM^ cells confirmed that mmu-miR-721 was released mainly by Th17^EAM^ cells (**Figure 1D**). A detailed characterization of the EVs secreted by both cell subsets showed that they barely differed among them, apart from the amount of EVs released which is greater in Treg^EAM^ cells-SN (**Figure S1D-F**). Moreover, mmu-miR-721 expression was significantly higher in EVs from the plasma of mice with EAM, compared to myocardial infarction and their respective controls (**Figure 1E**). Interestingly, such increased expression was not seen in mmu-miR-483-5p, underscoring mmu-miR-721 expression as a sensitive and specific of acute myocarditis in murine plasma.

**Figure 1.**
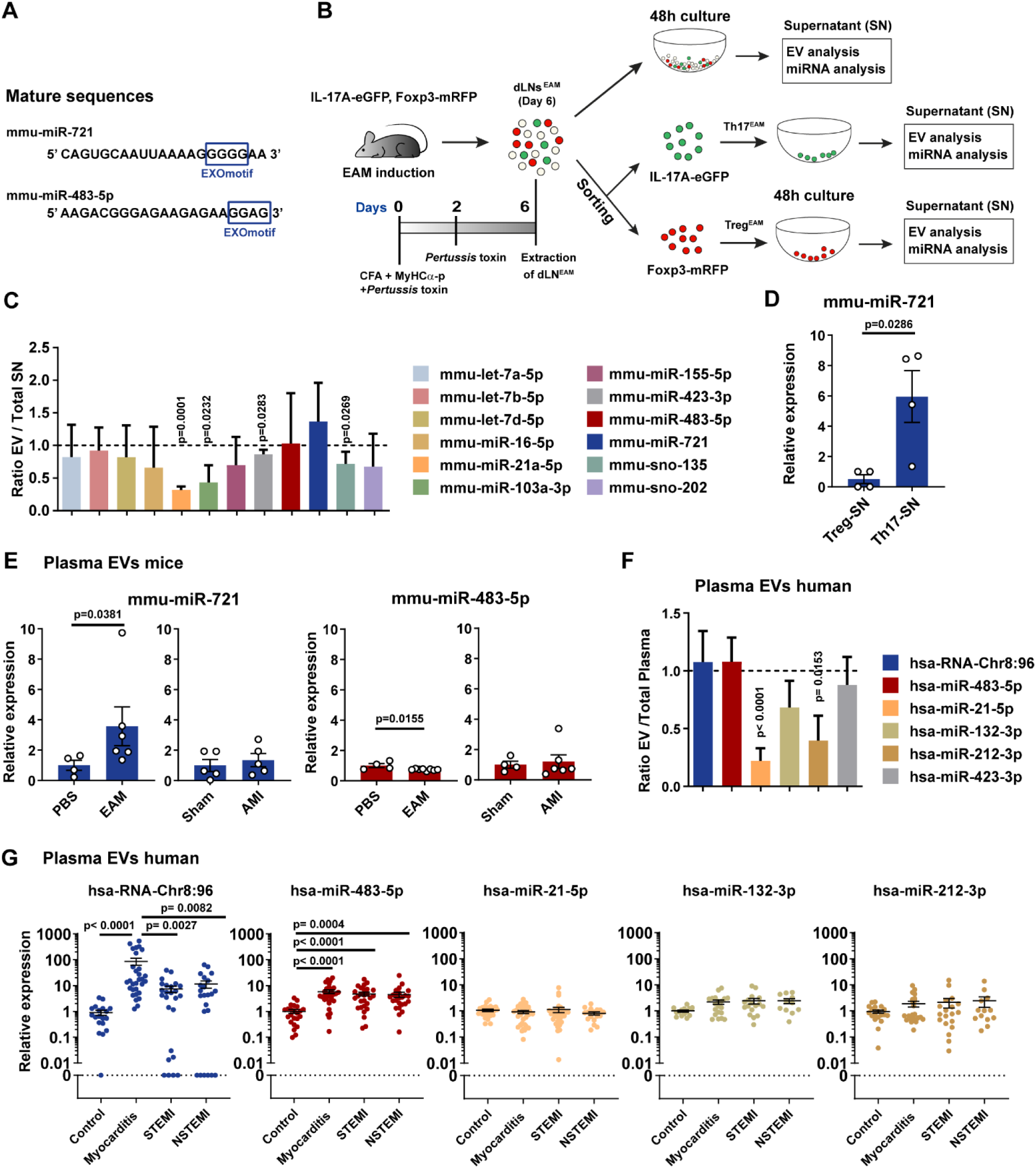
Th17-derived mmu-miR-721 and has-RNA-Chr8:96 is sorted into extracellular vesicles in mice and humans. (**A**) Mature sequences of mmu-miR-721 and mmu-miR-483-5p, putative EXOmotifs highlighted in blue. (**B**) Schematic representation of the protocol for extracellular vesicles (EV) analysis. Experimental autoimmune myocarditis (EAM) was induced in double reporter mice (IL-17-eGFP/Foxp3-mRFP) with MyHCα-p, CFA and *Pertussis* toxin. Draining lymph nodes (dLN; axillary, brachial and inguinal) were collected at day 6 post-induction and were stimulated *ex vivo* for 48h as the total cell pool (dLN^EAM^) or after Treg (Treg^EAM^) or Th17 (Th17^EAM^) sorting by FACS using Foxp3-mRFP^+^ and IL-17-eGFP^+^ markers respectively. EVs from cell culture supernatants were analysed. (**C**) Ratio between the relative expression of miRNAs in PEG-isolated EVs from supernatants (SN) *versus* total SN of dLN^EAM^, calculated after correction for the yield of the extraction using UniSp5 spike-in control (n=4-10). Dotted line indicates ratio =1. Data pooled from 3 independent experiments. Data is shown as means ± SEM and are analyzed against a hypothetical value 1 by one-sample *t*-test. (**D**) Mmu-miR-721 relative expression in PEG-isolated EVs from sorted Th17^EAM^- and Treg^EAM^-SN as shown in **A**. (n=4 of 6 pooled mice). Data are represented as mean ± SEM and were analyzed by Mann-Whitney U test. (**E**) Quantification of mmu-miR-721 and mmu-miR-483-5p by qPCR in PEG-isolated EVs from plasma of acute EAM (day 21) and acute myocardial infarction (AMI; day 3) mice, compared with respective controls (n=4-8). Data are represented as mean ± SEM and were analyzed by unpaired t-test or, when non-normal distributions, by Mann-Whitney U test for non-normal distributions. The ratio of relative miRNA expression in plasma PEG-isolated EVs *versus* total plasma from myocarditis patients (n=11-16). Data are represented as means ± SEM and analyzed against a hypothetical value 1 by one-sample *t*-test. (**G**) Quantification of miRNAs in PEG-isolated EV fractions from plasma of healthy donors (n= 25), myocarditis patients (n= 32), ST-segment elevation myocardial infarction (STEMI) patients (n= 35) and NSTEMI (non-STEMI) patients (n=24). Data are represented as means ± SEM of log10 of miRNA relative expression. Data were analyzed by one-way ANOVA with Tukey’s *post hoc* test or by Kruskal-Wallis with Dunn’s *post hoc* test when data are not normally distributed.

Additionally, the expression of selected hsa-miRNAs was studied in the EV-fraction after PEG precipitation from human plasma from 32 myocarditis patients. Whereas both hsa-RNA-Chr8:96 and hsa-miR-483-5p were sorted into the EV-fraction (**Figure 1F**), hsa-RNA-Chr8:96 detection was increased in plasma-EVs from acute myocarditis patients compared to STEMI, NSTEMI and healthy donors. hsa-miR-483-5p was increased in all patients with myocardial disease compared to healthy donors whereas hsa-miR-5p, hsa-miR-132-3p and hsa-miR-212-3p are unspecific of these populations (**Figure 1G**). Collectively, these data demonstrate that miR-721 is predominantly secreted in association with EVs and reaches the circulation encapsulated within EVs rather than as a freely soluble miRNA.

### Mmu-miR-721 inhibits *Pparg* enhancing *Rorc* and Th17 responses in EAM

We next investigated whether EV-associated miR-721 modulates Th17 cell biology and influences the development of EAM. The pre-mmu-miR-721 was overexpressed in dLN-EAM cells by lentiviral transduction. Lentiviral vectors encoding precursor mmu-miR-721 were designed (**Figure 2A**) and overexpression of the pre-miRNA confirmed by PCR amplification (**Figure 2B**) and the mature mmu-miR-721 by RT-qPCR (**Figure 2C**) in the dLN-EAM cells. Overexpression of mmu-miR-721 resulted in increased CD4^+^ IL-17A^+^ cell differentiation (**Figure 2D**) and enhancement of IL-17A production in the transduced cells at mRNA levels (**Figure 2E**) in dLN-EAM. It has been previously reported that *Pparg* is a direct target of miR-721^6^ and that this nuclear receptor negatively regulates Th17 cell differentiation by controlling the orphan nuclear receptor *Rorc* induction^8^, a master transcription factor of Th17 cells. Hence, to understand how mmu-miR-721 regulates Th17 cell function, we analyzed the expression of *Pparg* and *Rorc* at the mRNA level. We confirmed a reduction in *Pparg* transcription after mmu-miR-721 overexpression in dLN-EAM cells (**Figure 2E**). Conversely, we observed increased *Rorc* transcription together with enhanced RORγt expression in CD4^+^ T cells from dLN-EAM (**Figures 2E and 2F**), supporting a role for mmu-miR-721 in promoting Th17 responses during myocarditis development through regulation of the *Pparg* / *Rorc* axis.

**Figure 2.**
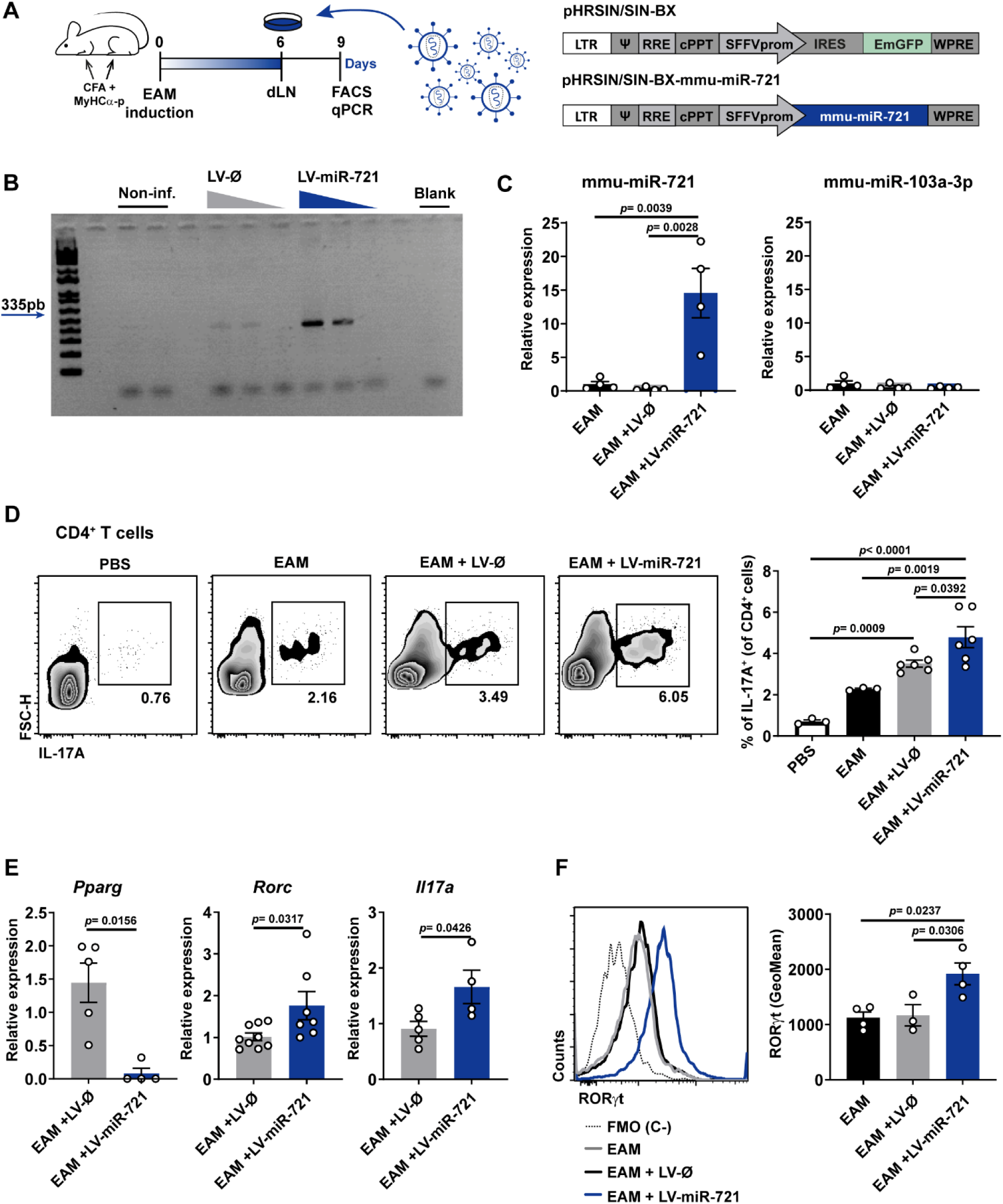
Overexpression of mmu-miR-721 *ex vivo* enhances Th17 responses. **(A**) Schematic representation of the experimental setup and the lentiviral vector generated for the overexpression of mmu-miR-721 in dLN^EAM^-derived cells. LTR (5′long terminal repeat); Sequence Ψ (packaging sequence); RRE (*Rev* response element); cPPT (central sequence Polypurine Tract); SFFV (lentiviral promoter); IRES (internal ribosome entry site); WPRE (Woodchuck hepatitis virus (WHP) Post-transcriptional regulatory element); mmu-miR-721 (mmu-pre-miR-721-encoding sequence); EmGFP (Emerald Green Fluorescent Protein-encoding sequence). (**B**) Analysis by PCR of mmu-miR-721 expression in dLN^EAM^ cells after transduction with lentiviral vectors (LV)-empty (Ø) or -miR-721 MOI 5, MOI 2 and MOI 1. (**C**) Quantitative analysis of mmu-miR-721 and control mmu-miR-103a expression after transduction with MOI 5 in dLN^EAM^ cells. Rnu1a1 small nuclear RNA was used as normalizer for qPCR. Data are analyzed with one-way ANOVA and Tukey’s *post hoc* test. (**D**) FACS analysis of Th17 cells in dLN^EAM^ cells or control vaccination with PBS, and transduction or not with LV-empty or LV-miR-721 (MOI 5). Data is analyzed by one-way ANOVA with Holm-Sidak’s *post hoc* test. (**E**) qPCR analysis of *Pparg*, *Rorc* and *Il17a* genes in dLN^EAM^ cells after LV-empty or LV-miR721 transduction. Data were analyzed with unpaired *t*-test and Mann-Whitney *U* test for non-normal distributions. (**F**) RORγt expression in dLN^EAM^ cells analyzed by FACS. Dashed line indicates fluorescence minus one (FMO) negative control. One-way ANOVA with Tukey’s *post hoc* test. Data in **B-F** correspond to one representative out of three independent experiments (n=3-9 per group) and are represented as means ± SEM.

To assess whether the increased Th17 response is a direct result of *Pparg* downregulation and test the potential interference with other signaling axes, we evaluated these parameters in a mouse model of inducible *Pparg* depletion in the hematopoietic compartment (Mx^WT/*Cre*^ *Pparg^fl/fl^*). We analyzed the effect of mmu-miR-721 overexpression in CD4^+^ T cells from dLN-EAM by FACS. RORγt and IL-17A were efficiently induced in Mx*^Cre^ Pparg^fl/fl^* in comparison to Mx^WT^ *Pparg^fl/fl^* mice-derived CD4^+^ T cells (**Figure 3A**). However, while mmu-miR-721 overexpression did not have any further impact on RORγt expression, it enhanced IL-17 expression in Th17 cells, suggesting a PPARγ-independent pathway contributing to this increase (**Figure 3A**). In parallel, we confirmed that both *Rorc* and *Il17a* mRNA levels showed the same expression pattern as those observed in FACS analysis, proving that the differences occur at a transcriptional level (**Figure 3B**). Altogether, these results demonstrate that the effect of miR-721 on RORγt expression is mediated by *Pparg* inhibition.

**Figure 3.**
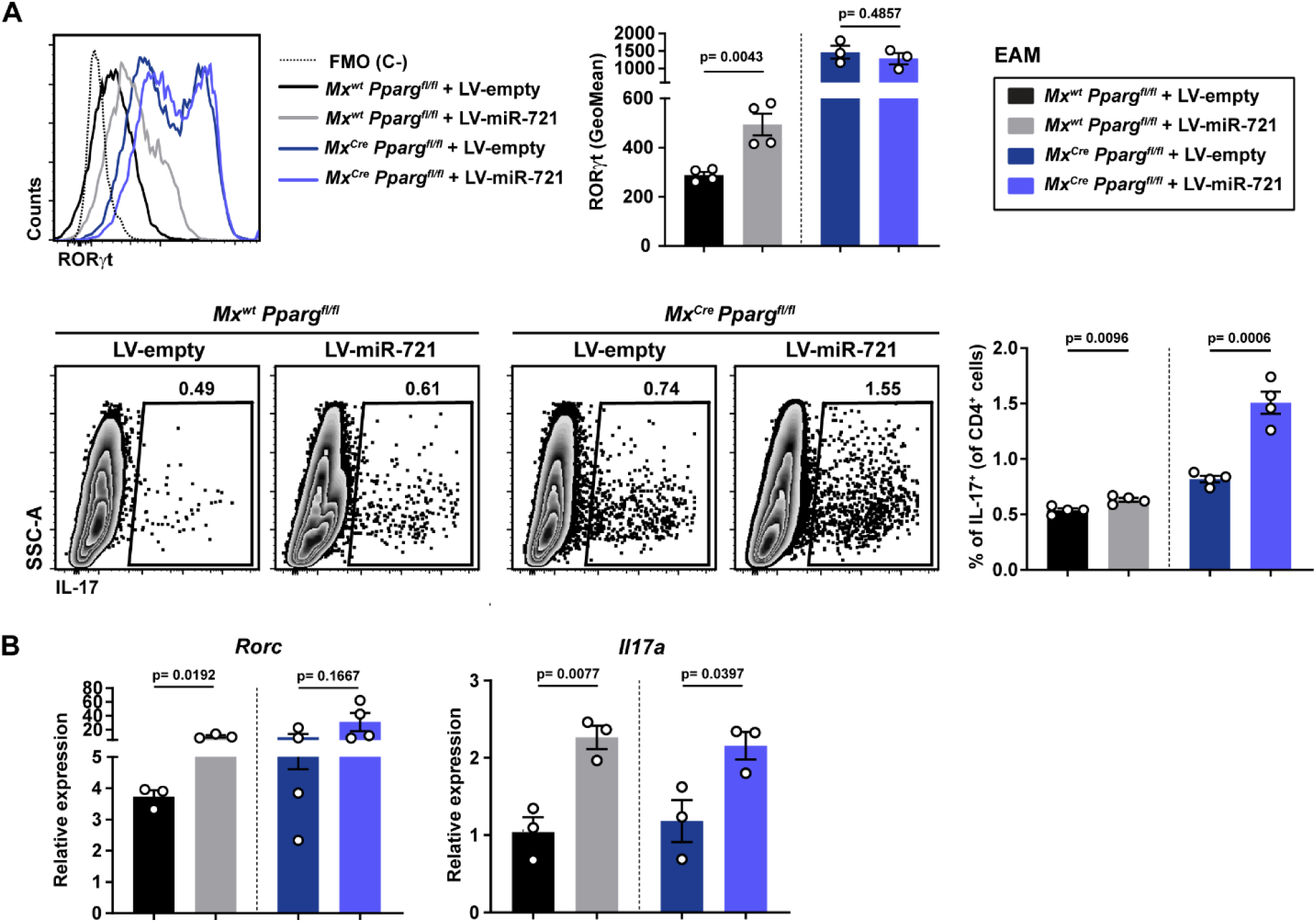
Ablation of *Pparg* prevents miR-721-mediated RORγt upregulation. **(A**) FACS quantification of RORγt expression in CD4^+^ T cells and percentages of Th17 cells in dLN^EAM^-derived cells from Mx^wt^ or Mx^Cre^ Pparg^fl/fl^ mice after lentiviral transduction with LV-empty or LV-miR-721 (MOI 5). FMO (fluorescence minus one) indicates negative staining control. (**B**) Analysis of *Rorc* and *Il17a* mRNA levels by qPCR. Data in **A** and **B** are correspond to one representative out of three independent experiments (n=3-5 per group). Data are shown as mean ± SEM and were analyzed with unpaired *t*-test and Mann-Whitney *U* test for non-normal distributions.

### Human homolog hsa-RNA-Chr8:96 targets *PPARG* mRNA expression

After confirming that hsa-RNA-Chr8:96 is also sorted into EVs and learning about the murine miR-721 influence in Th17 responses, we addressed whether the human homolog can control Th17 via the *Pparg/Rorc* axis. No gene has so far been validated as a target of hsa-RNA-Chr8:96 action. To address this, we cloned the 3’-UTR region of *PPARG* in a luciferase reporter vector system, which was co-transfected with an empty expression vector or one encoding hsa-RNA-Chr8:96 (**Figure 4A**) to test whether the miRNA can physically bind to its putative target site (**Figure S2**). The luciferase assay revealed decreased luminescence signal when the miRNA was expressed (**Figure 4B**), indicating hsa-RNA-Chr8:96 direct binding to *PPARG* 3’-UTR region. Further, to determine the consequences of this binding on gene expression we transduced primary human peripheral blood lymphocytes with lentiviral vectors carrying or not hsa-RNA-Chr8:96 sequence. Upon miRNA overexpression, levels of *PPARG* mRNA were significantly reduced (**Figure 4C**).

**Figure 4.**
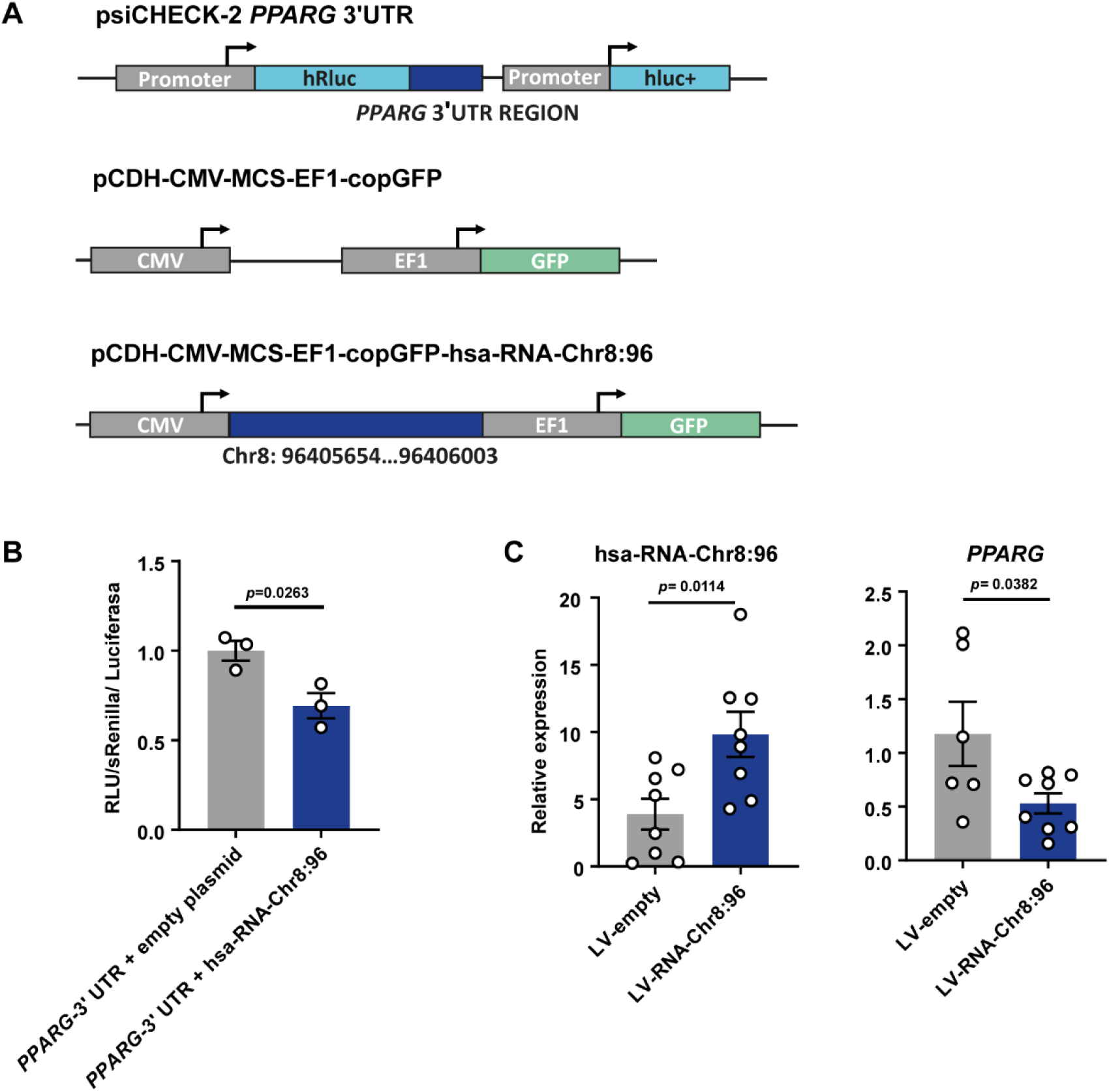
Validation of *PPARG* as a target gene of hsa-RNA-Chr8:96. (**A**) Schematic representation of the plasmids used for transfection of HEK-293 cells in **B** or for generation of LVs used for transduction of isolated human peripheral blood lymphocytes in **C**. *hRluc* (*Renilla* luciferase); *hluc*+ (Firefly luciferase); CMV (cytomegalovirus promoter); EF1 (elongation factor 1 alpha); GFP (Green Fluorescence Protein). (**B**) Luciferase reporter assay in HEK-293 cells showing luminescence activity after co-transfection of psiCHECK-2 construct encoding *PPARG* 3’-UTR region with either an expression plasmid empty or encoding hsa-RNA-Chr8:96 sequence. Luminescence values of *Renilla* luciferase were normalized by firefly luciferase and by the values obtained with cells transfected with the empty construct (n=3). (**C**) Quantification of hsa-RNA-Chr8:96 and *PPARG* mRNA expression in isolated human peripheral blood lymphocytes after transduction with the lentiviral vector containing the hsa-RNA-Chr8:96-encoding plasmid (LV-miR-Chr8:96) or the control plasmid (LV-empty), analysed by qPCR (n=6-8). Data pooled from 2 out of 3 independent experiments. Data is represented as mean ± SEM and were analyzed by unpaired *t*-test.

### Inhibition of mmu-miR-721 ameliorates EAM development *in vivo*

Since mmu-miR-721 controls myocarditis-induced Th17 responses, we next aimed to ascertain whether the functional blockade of this miRNA has an implication in the progression of the disease. For this purpose, we designed a miRNA sponge molecule with 4 tandem-repeated reverse complementary sequences of the mature mmu-miR-721 separated by spacers, namely, random sequences, that is able to block and inhibit the miRNA (miR-721-sponge, **Figure 5A**). EAM was induced in BALB/c by subcutaneous immunization with the MyHCα−p along with either the miR-721-sponge or a scramble sequence of similar size as a control. Analysis by qPCR confirmed mmu-miR-721 downregulation (**Figure 5B**) and overexpression of its target *Pparg* (**Figure 5C**) in dLN-EAM from the sites of injection in the presence of the miR-721-sponge 2 days after its delivery.

**Figure 5.**
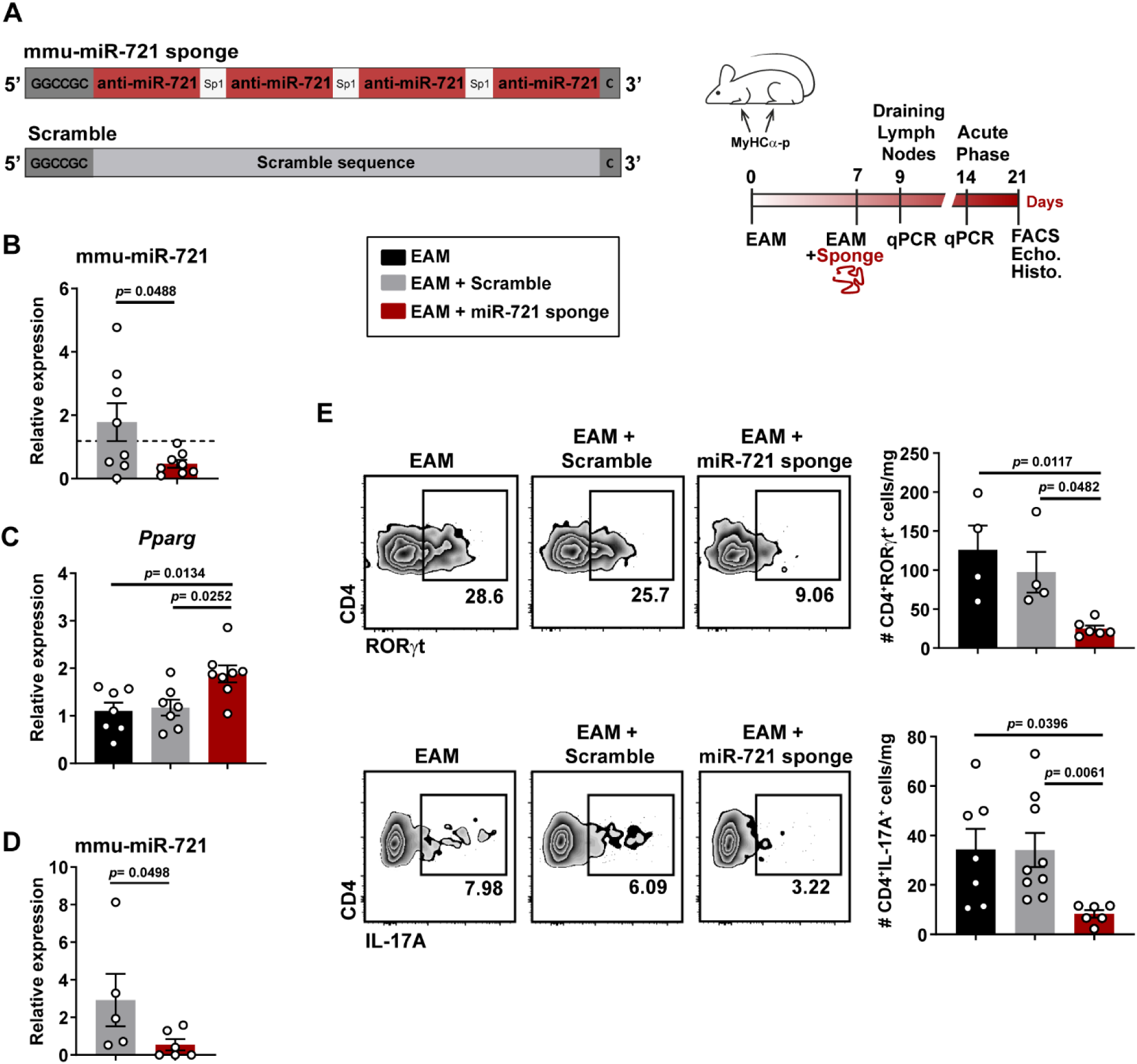
Inhibition of mmu-miR-721 allows PPARγ mediated repression of Th17 responses *in vivo*. (**A**) Schematic view of mmu-miR-721 sponge molecule (miR-721-sponge) for silencing mmu-miR-721 and experimental setup of the disease and harvest timepoints. Anti-miR-721, reverse complementary sequence of the mature mmu-miR-721; SP1, spacer sequences of 4-6 nt. A scramble non-complementary nucleotide sequence of the same size was used as control. (**B**) Blocking efficiency of miR-721 and (**C**) relative expression of *Pparg* mRNA in dLN at day 9 after EAM induction (n=7-8), analyzed by qPCR. **B** and **C** show data pooled from two independent experiments. In **B**, dotted line represents the levels of the miRNA in EAM mice. (**D**) Expression of miR-721 in circulating plasma at day 14. (n= 5-6) (**E**) Analysis by FACS of percentages of RORγt+ and IL-17+ heart-infiltrating CD4+ T cells 21 days after EAM. Representatives density plots and cell numbers per mg of heart tissue are shown (n= 4-9). Data are represented as means ± SEM. One representative out of four independent experiments. Data in **B** and **D** were analyzed by unpaired *t*-test. In **C** and **E**, one-way ANOVA with Holm-Sidak’s *post-hoc* test or Kruskal-Wallis with Dunn’s *post-hoc* test when non-normal distributions.

miR-721 blockade was further effective in circulation too as its levels in plasma appeared downregulated at the start of the acute phase at day 14 (**Figure 5D**). Recruitment of heart-infiltrating CD4^+^ T cells expressing RORγt and IL-17A is significantly inhibited in mice treated the miR-721-sponge (**Figure 5E**), assessed twenty-one days after EAM induction, during the peak of the acute phase of myocarditis and supporting the role of miR-721 controlling Th17 responses. To study the implications of miR-721 inhibition on disease progression, physical, functional and inflammatory parameters were taken at day 21 after EAM induction. Mice treated with the miR-721-sponge exhibit reduced heart inflammation, measured by decreased number of heart-infiltrating leukocytes (**Figure 6A**). Infiltrating myeloid cells and, more specifically, macrophages were reduced, marked by their surface expression of CD11b and F4/80 (**Figure 6B**) and known to promote cardiac damage^15^. Decreased inflammation is accompanied by improved heart-to-body weight and heart-to-tibia length ratios (**Figure 6C**) together with reduced left ventricular mass analyzed by echocardiography (**Figure 6D**). Additionally, left ventricular function was conserved (**Figure 6E**), suggesting an overall reduction in myocardial disease. Thus, the functional blockade of mmu-miR-721 protects against the development of myocarditis by inhibiting Th17 responses.

**Figure 6.**
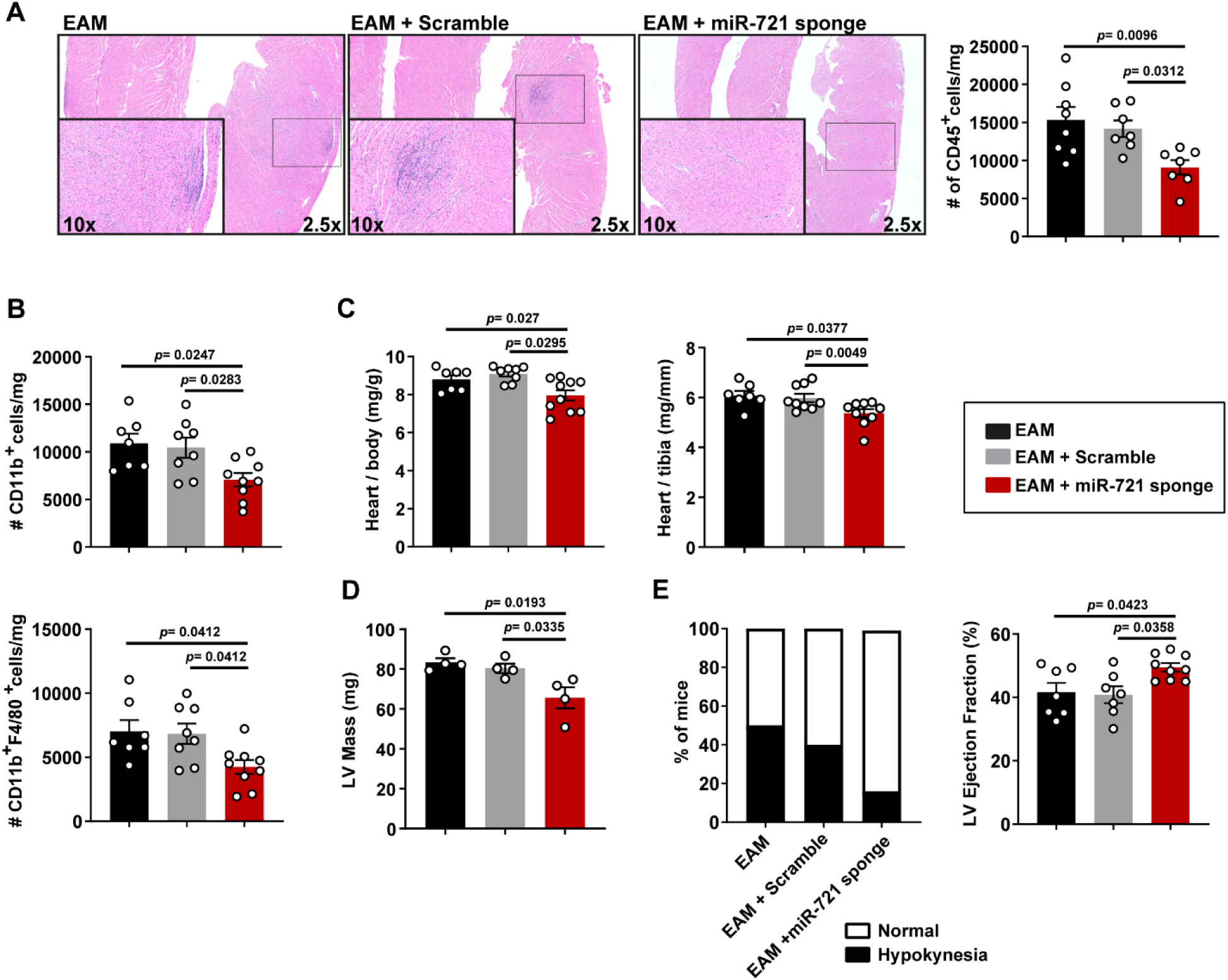
Blockade of mmu-miR-721 ameliorates EAM development *in vivo*. (**A**) Representative hematoxylin eosin staining of heart sections 21 days after EAM induction. Cell infiltrate indicates the number of heart-infiltrating CD45+ leukocytes per mg of heart (n= 4-6). (**B**) Number of CD11b+ and CD11b+F4/80+ cells per mg of heart. (n=7-9) (**C**) Ratio of heart weight (mg) to body weight (g) and to tibia length (mm) 21 days after EAM (n=7-9) (**D**) Left ventricle (LV) mass, assessed by transthoracic echocardiography (n=4) (**E**) Percentage of hypokinetic mice and LV ejection fraction as a measure of heart function, assessed by transthoracic echocardiography (n=4-9). Data are represented as means ± SEM. One representative out of four independent experiments. Data are analyzed by one-way ANOVA with Holm-Sidak’s *post-hoc* test.

## Discussion

Myocarditis remains a diagnostic challenge in clinical practice due to the atypical nature of its clinical presentation and the heterogeneity of its aetiology^1^. We have previously described the potential of mmu-miR-721 and hsa-RNA-Chr8:96 as diagnostic tools for the detection of acute myocarditis, regardless of aetiology and opposed to other miRNAs^4^. Previous studies have detected mmu-miR-721 presence in EVs in the SNC or SNC-related diseases^16,17^. In line with these, our findings confirm that both murine and human forms of this miRNA are enriched in EVs during EAM, contributing to its stability in plasma and enhancing its sensitivity as a liquid biopsy biomarker, protecting it from RNase degradation and enhancing its diagnostic value^18,19^. Furthermore, the specificity of other potential candidates such as miR-483-5p^4^ is proven again insufficient to discriminate against other cardiovascular pathologies despite its encapsulation and protection into EVs.

Aside from its potential as a tool, we have identified that this miRNA plays a role in the regulation of Th17 responses during acute myocarditis. Our data demonstrate that mmu-miR-721 overexpression downregulates *Pparg* in *ex vivo* dLN-derived cells isolated from EAM mice, revalidating this target in T cells. PPARγ is a repressor of *Rorc* transcription^8^. Consequently, mmu-miR-721 overexpression results in increased *Rorc* mRNA and RORγt protein levels, enhancing IL-17 production.

Remarkably, while RORγt levels experience no significant changes upon miR-721 overexpression in the absence of *Pparg*, IL-17 levels are increased. This suggest RORγt upregulation is dependant on PPARγ, but other molecular pathways might be playing a role in IL-17 expression. *Nos2* is one of the few target genes described for miR-721 activity^20^, but its role on Th17 cells is more controversial. Although NOS2-mediated generation of NO has been described to maintain stability of human Th17 cells^21^, NO functions as a brake for Th17 cell responses and cytokine production in different mouse models of autoimmunity by interfering with RORγt-mediated activation of *Il17a* promoter activation^22,23^. Thus, exploring the role of this enzyme within this regulatory axis could become important to target miR-721 in inflammatory contexts.

Inversely, our data show that the *in vivo* inhibition of mmu-miR-721 with blocking antisense sponge molecules augments *Pparg* levels in the dLN of EAM mice. This blockade of mmu-miR-721 decreases the amount of Th17 cells in the myocardium and ameliorates EAM progression in both physical and functional terms. Therefore, mmu-miR-721 works as an immunomodulator of myocarditis development by boosting Th17 responses and, more particularly, IL-17 production. Given that miR-721 is secreted in EVs, this feature could facilitate the amplification of Th17 responses during myocarditis, as Th17 cells differentiate in mediastinal lymph nodes and migrate to the myocardium where IL-17 promotes inflammation by inducing structural and immune cells to produce neutrophil-attracting chemokines such as CXCL1, CXCL2, CXCL5 and CXCL8 (IL-8), and by stimulating granulopoiesis through G-CSF^24–26^. IL-17 is at the core of disease development as it has been shown pivotal for B cell production of cardiac-specific antibodies^27^, recruitment of pro-inflammatory monocytes and macrophages^15^ and cardiac remodelling, fibrosis and progression to dilated cardiomyopathy (DCM)^26,28,29^. In this regard, this miRNA appears as a suitable candidate for the development of therapeutic approaches for myocarditis.

Our data also demonstrate that human PPARγ expression is targeted by hsa-RNA-Chr8:96, thereby offering the opportunity to translate the knowledge of this regulatory axis to the clinics. Targeting this molecule using anti-miRNA complexes as a therapeutic tool could either mitigate myocarditis development and/or reduce the risk of progression towards DCM. Further, the potential of controlling Th17 responses with this tool extends beyond myocarditis, offering new avenues for exploring similar therapeutic strategies to tackle other autoimmune and non-autoimmune disorders where Th17 cells play a detrimental role.

The active secretion of this miRNA into circulating EVs leads us to postulate that it may play a role not only within Th17 cells but also in distant cells and tissues. Other known contributors of IL-17 secretion such as γδT cells, CD8 T cells or some innate cells^30^ could also be targets of Th17-derived miR-721 paracrine action. Specific experiments designed to identify the target cells that uptake the Th17-cell-derived EV-mmu-miR-721 in myocarditis would clarify a possible function of this miRNA in intercellular communication in such context.

Overall, our findings expand the diagnostic potential of miR-721 for the detection of acute myocarditis and presents this miRNA as a therapeutic target to reduce pro-inflammatory Th17 responses, myocarditis burden and subsequent development of heart failure.

## Acknowledgements

We thank the patients and other volunteers for their participation in this study; Ana Vanesa Alonso and Lorena Flores for their excellent technical work with the echocardiography acquisition and analysis in the murine models; Ramón F. Maruri for his work analyzing data for the patients from Hospital de la Princesa; Laura Fernández and Belen Díaz from HM Hospitales for patient recruitment; and the staff members at the Genomics, Advanced Imaging, Cellomics, and Comparative Medicine units at the CNIC for their support in this work.

## Tables

**Supplementary Table 1.**
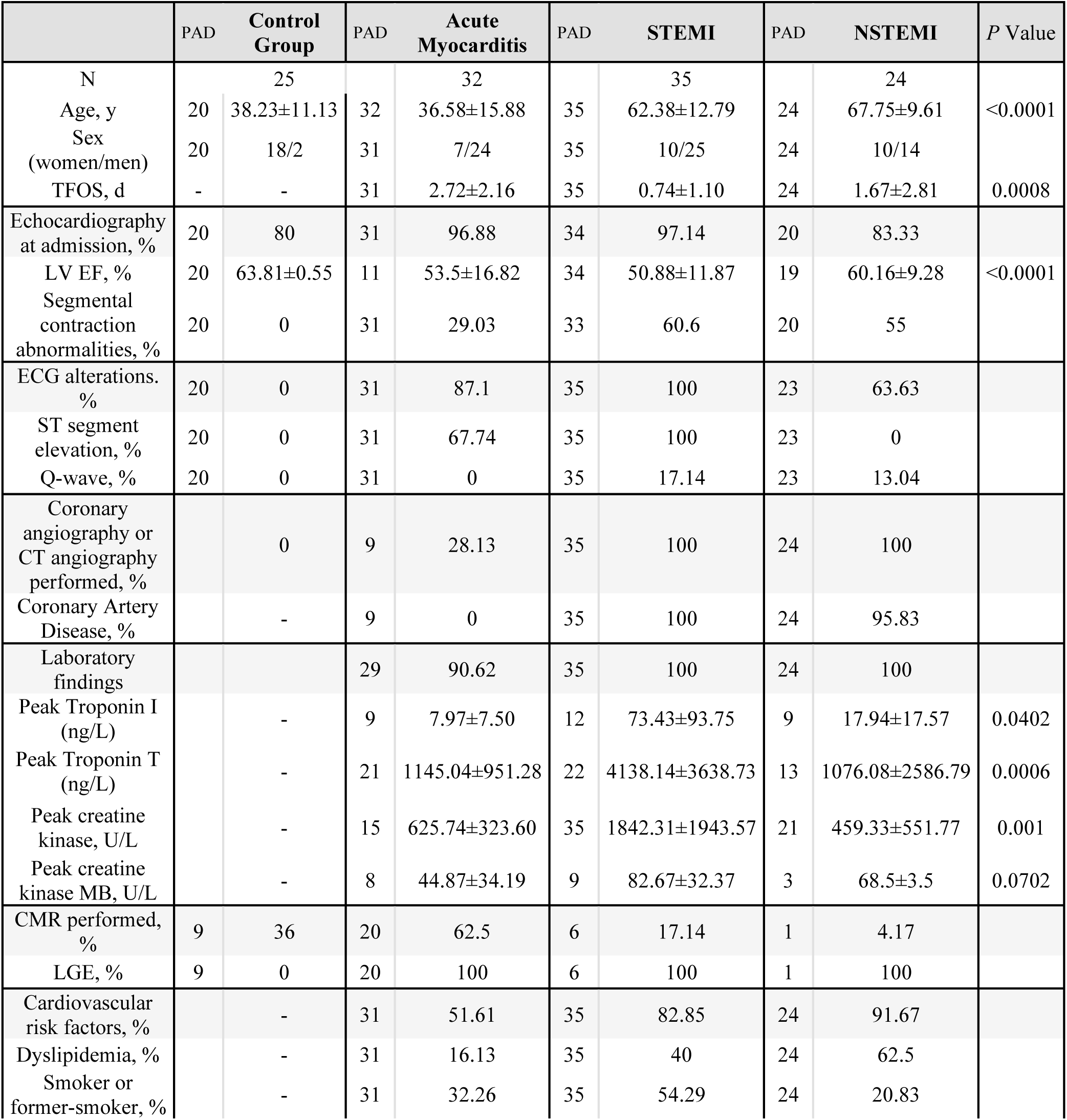
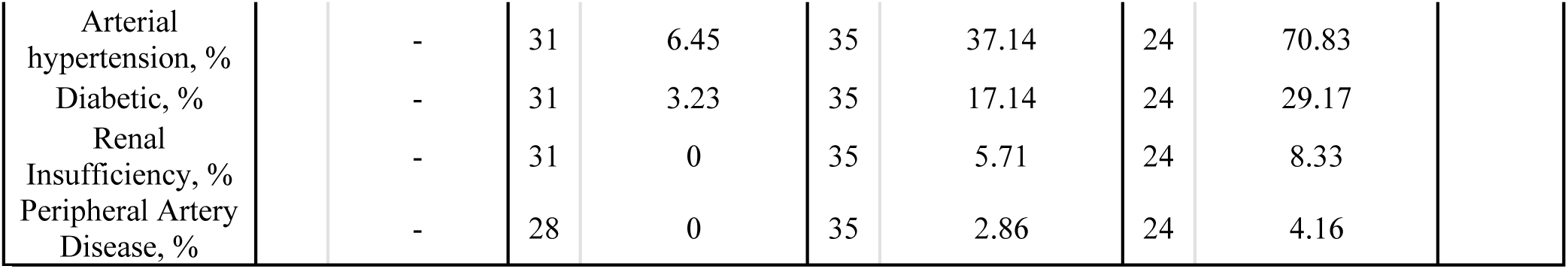
Baseline characteristic of the Study Population. Data are expressed as means ± SD or as percentages of patients. PAD, patients with available data; TFOS, time from the onset of symptoms; LV EF, left ventricle ejection fraction; CMR, cardiovascular magnetic resonance; LGE, late gadolinium enhancement. P-value calculated by one-way ANOVA.

---

**Figure S1.**
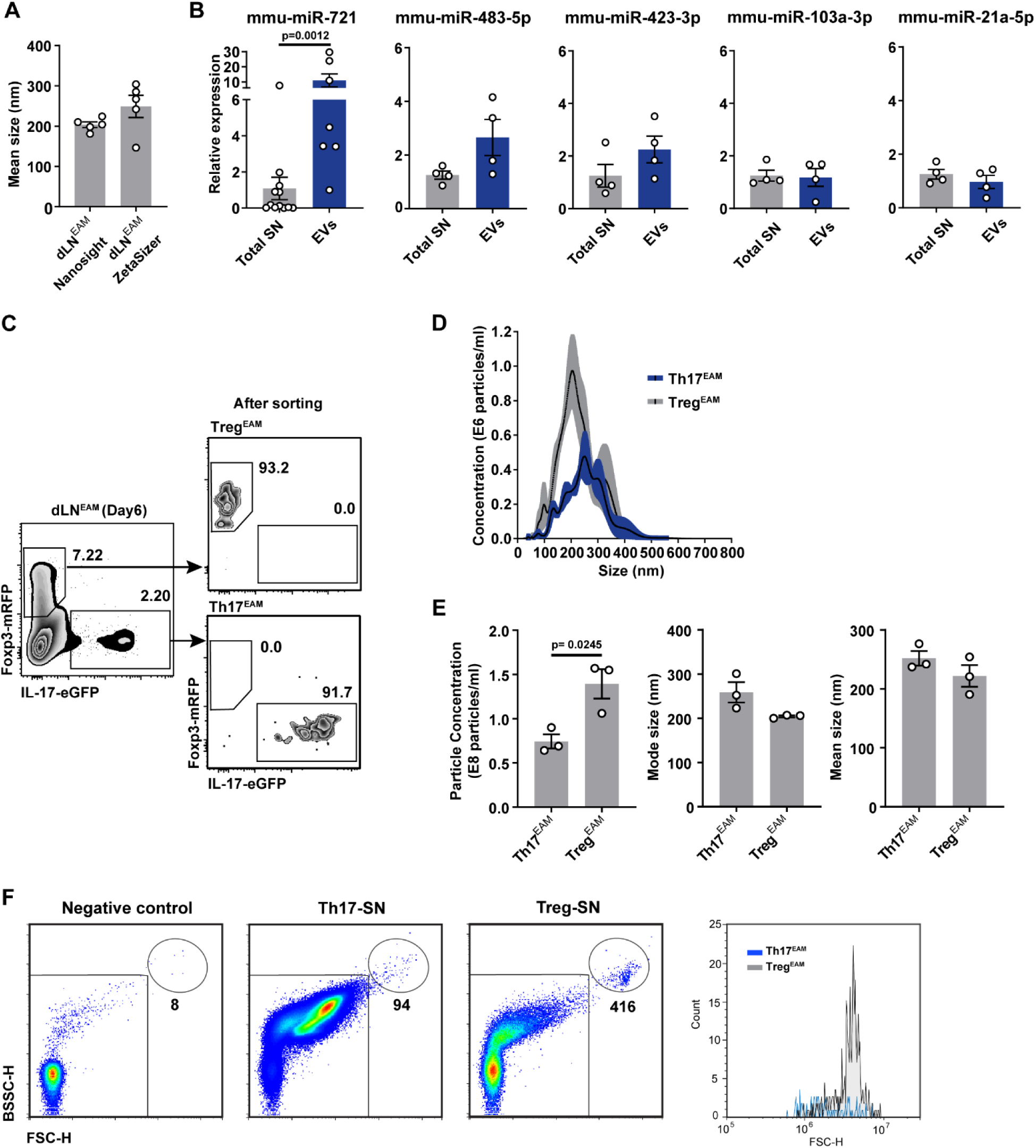
Characterization of T-cell-derived EVs from EAM mice. (**A**) Mean size of PEG-isolated EVs from SN of dLN^EAM^-derived cells cultured for 48h and analyzed by *Nanosight* and *ZetaSizer* (n=5). (**B**) Relative expression of miRNAs in PEG-isolated EVs from SN *versus* total SN, after evaluating the most stable normalizers between both conditions (n=4-10). (**C**) Representative dot plot of the percentages of Foxp3-mRFP^+^ and IL-17-eGFP^+^ cells (gated on CD4^+^ cells) in dLN^EAM^. (**D**) Size distribution of EVs from sorted Th17^EAM^- and Treg^EAM^-SN, analyzed by *Nanosight*. Black lines in histogram indicates the mean and shadows represent the SEM. (**E**) Quantification of equivalent particle concentration and size (mean/mode) of EVs derived from sorted Th17^EAM^ and Treg^EAM^ by *Nanosight* (n=3 of 4-6 pooled mice). (**F**) Visualization and quantification of EV particles from Th17^EAM^- and Treg^EAM^-SN by flow cytometry. Negative control corresponds to media. Data in **A**, **B** and **E** are expressed as mean ± SEM and were analyzed by Unpaired *t*-test or Mann-Whitney U-test for non-normal distribution. Data in **A** and **E** correspond to one representative out of 3 independent experiments.

**Figure S2.**
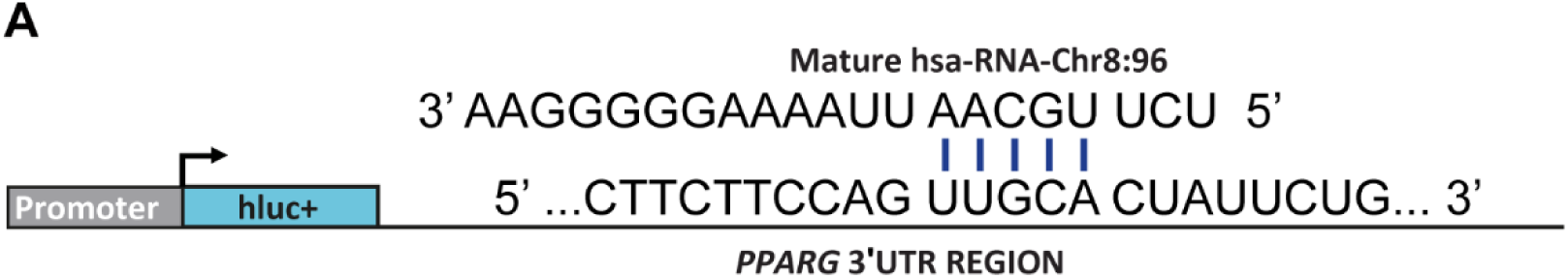
miR-721 binding site in *PPARG* 3’-UTR. (**A**) Schematic representation of the proposed binding site for the mature miR-721 sequence within *PPARG* 3’-UTR.

